# LDER-GE estimates phenotypic variance component of gene-environment interactions in human complex traits accurately with GE interaction summary statistics and full LD information

**DOI:** 10.1101/2023.11.22.568329

**Authors:** Zihan Dong, Wei Jiang, Hongyu Li, Andrew T. DeWan, Hongyu Zhao

## Abstract

Gene-environment (GE) interactions are essential in understanding human complex traits. Identifying these interactions is necessary for deciphering the biological basis of such traits. In this study, we introduce a statistical method Linkage-Disequilibrium Eigenvalue Regression for Gene-Environment interactions (LDER-GE). LDER-GE improves the accuracy of estimating the phenotypic variance component explained by genome-wide GE interactions using large-scale biobank association summary statistics. LDER-GE leverages the complete Linkage Disequilibrium (LD) matrix, as opposed to only the diagonal squared LD matrix utilized by LDSC (Linkage Disequilibrium Score)-based methods. Our extensive simulation studies demonstrate that LDER-GE performs better than LDSC-based approaches by enhancing statistical efficiency by approximately 23%. This improvement is equivalent to a sample size increase of around 51%. Additionally, LDER-GE effectively controls type-I error rate and produces unbiased results. We conducted an analysis using UK Biobank data, comprising 307,259 unrelated European-Ancestry subjects and 966,766 variants, across 151 environmental covariate-phenotype (E-Y) pairs. LDER-GE identified 35 significant E-Y pairs while LDSC-based method only identified 25 significant E-Y pairs with 23 overlapped with LDER-GE. Furthermore, we employed LDER-GE to estimate the aggregated variance component attributed to multiple GE interactions, leading to an increase in the explained phenotypic variance with GE interactions compared to considering main genetic effects only. Our results suggest the importance of impacts of GE interactions on human complex traits.

## Introduction

A growing body of literature underscores the significant role of gene-environment (GE) interactions in shaping human complex traits^1-4^. The exploration of GE interactions may elucidate a portion of the ‘missing heritability’^5^ — the phenotypic variance not accounted for by known genetic effects. Additionally, the inference of GE interactions and their effects can contribute to our understanding of human disease etiology and mechanisms^6^, and enhance our ability to assess risk and identify high-risk individuals, ultimately supporting the development of personalized medicine^1^. Traditionally, environmental exposure variables have been limited to factors like environmental toxins, air pollutants, or viral infections^6^. However, some gene-environment interaction studies^7,8^ also consider other variables, heritable or non-heritable, such as sex, as environmental exposure variables. In this study, we adopt a broad definition, considering both non-heritable covariates and heritable phenotypes as environment interactive variables, as previously discussed^2^.

Numerous methods and tools have been developed to investigate GE interactions from various angles. One such approach is the genome-wide interaction scan (GWIS), which estimates the interaction effect^9^ between individual genetic variants and environmental factors through regression. GWIS generates interaction summary statistics for each variant, akin to conventional genome-wide association studies (GWAS). However, we note that GE interaction effect sizes tend to be smaller than genetic main effects^10^. Consequently, this can lead to reduced statistical power, particularly when challenged by the multiple testing burden across the entire genome^11^. Several studies have directed their efforts towards estimating the genome-wide contribution of GE interactions through diverse statistical approaches. One such method is the Gene-Environment Interaction Genomic Restricted Maximum Likelihood (GEI-GREML), which leverages restricted maximum likelihood estimation by pre-computing the correlation matrix of the GE term across samples^12^. On the other hand, the Multivariate Reaction Norm Model (MRNM) is a reaction norm model that has the capability to distinguish between GE interaction and GE correlation^13^. Both GEI-GREML and MRNM necessitate individual-level genotype data and can be computationally demanding and time-consuming, especially when dealing with extensive biobank datasets.

To tackle these challenges, researchers have devised alternative methods that make use of GWIS summary statistics. Notably, methods like PIGEON^7^ and GxEsum^8^ build upon the principles of LD-score regression (LDSC)^14^. They harness partial linkage disequilibrium (LD) information among genetic variants to estimate the phenotypic impact of GE interactions using the method of moments. However, this approach often results in reduced statistical efficiency when estimating variance components, because the phenotypic variance attributed to GE interactions is often considerably smaller than the narrow-sense heritability. For example, across a dataset encompassing more than 500 traits, the phenotypic variance explained by genetic-sex interactions typically falls within the range of 0% to a maximum of 2%^7^. While this may appear modest, acknowledging and investigating this component remains important for our understanding of complex traits and disease etiology. An inefficient estimation method may fail to detect the contribution of GE interactions. Consequently, there is a need for a more efficient approach to estimate the phenotypic variance explained by GE interactions while effectively managing computational demands. Current LDSC-based frameworks^7,8,14^ make use of the squared variant LD matrix but primarily focus on diagonal information. Previous research^15,16^ has convincingly shown that incorporating the complete LD information can substantially enhance the efficiency of estimating narrow-sense heritability under the genetic additive effect model. Building upon this insight, we introduce the Linkage-Disequilibrium Eigenvalue Regression for Gene-environment interactions (LDER-GE) to estimate the genome-level GE interaction variance component more efficiently.

LDER-GE mimics the original LDER framework^15^, which harnesses the full potential of LD information through eigen-decomposition of the LD matrix. This process transforms the original GWIS summary statistics and consolidates the association information. Notably, LDER-GE relies on summary statistics and the LD matrix constructed using a reference panel. Consequently, it efficiently manages large-scale Biobank data without imposing substantial computational demand. Extensive simulations provide evidence that both LDER-GE and the LDSC-based method effectively control the type-I error rate and deliver unbiased estimates. However, LDER-GE excels in terms of statistical efficiency compared to the LDSC-based method in all simulation scenarios. In a real-data application involving 151 E-Y pairs from the UK-Biobank^17^, the LDSC-based method identified 25 GE interaction signals, whereas LDER-GE identified 35 E-Y pairs (40% increase). For a more precise assessment of the contribution of GE interactions to missing heritability, we estimate the aggregated GE interaction variance involving multiple environmental covariates and the analyzed phenotypes. In this regard, LDER-GE facilitates more accurate estimation. In summary, the missing heritability contributed by the aggregated multi-covariate GE interaction variance represents a substantial addition to the narrow-sense heritability.

## Results

### Method overview

We propose LDER-GE to improve statistical efficiency with summary-level association statistics. Under the polygenic GE model where each standardized variant-by-E term has small effect, the expectation of cross-variant level GE interaction association chi-square statistics is (details in methods and supplementary note 1)

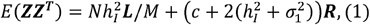

where **Z** is the original GWIS Z-score vector, **R** is the LD matrix, **L = R**^T^**R** is the LD score matrix, N is the sample size of the GWAS summary statistics, c is the unconstrained intercept with potential inflation,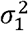 is the non-genetic environment interaction variance and 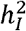 is the explained variance of the GE interaction. We incorporate full LD information by conducting eigen-decompose the LD matrix as **R** = **UDU**^T^, with **U** being the orthogonal matrix of eigenvectors and **D** being the diagonal eigen value matrix, and transforming the original GWIS Z-score vector 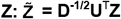, yielding

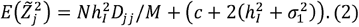

We run iterative weighted least squares to estimate 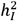 and the intercept, followed by delete-block-wise jackknife to estimate the standard error.

### Simulation results using real genotype panel

Figure 1 and Table 1 compare the performance of LDER-GE and the LDSC-based method across various parameter combinations. Across all simulation scenarios, the LDER-GE method consistently had better performance than the LDSC-based method, whether utilizing an UKBB in-sample or 1000 Genomes project out-sample reference panel. We assessed performance using precision, which is the inverse of the standard deviation, and root mean squared error. Notably, this precision improvement was, on average, equivalent to a 51% increase in sample size when analyzing continuous simulated phenotypes during in-sample estimation. Table 2 shows that the type-I error rates were well-controlled for all three methods: LDER-GE using an in-sample LD panel, LDER-GE using an out-sample LD panel, and LDSC-based in-sample LD panel. This held true both in scenarios with and without the presence of non-genetic residual-environment interaction effects.

**Figure 1:**
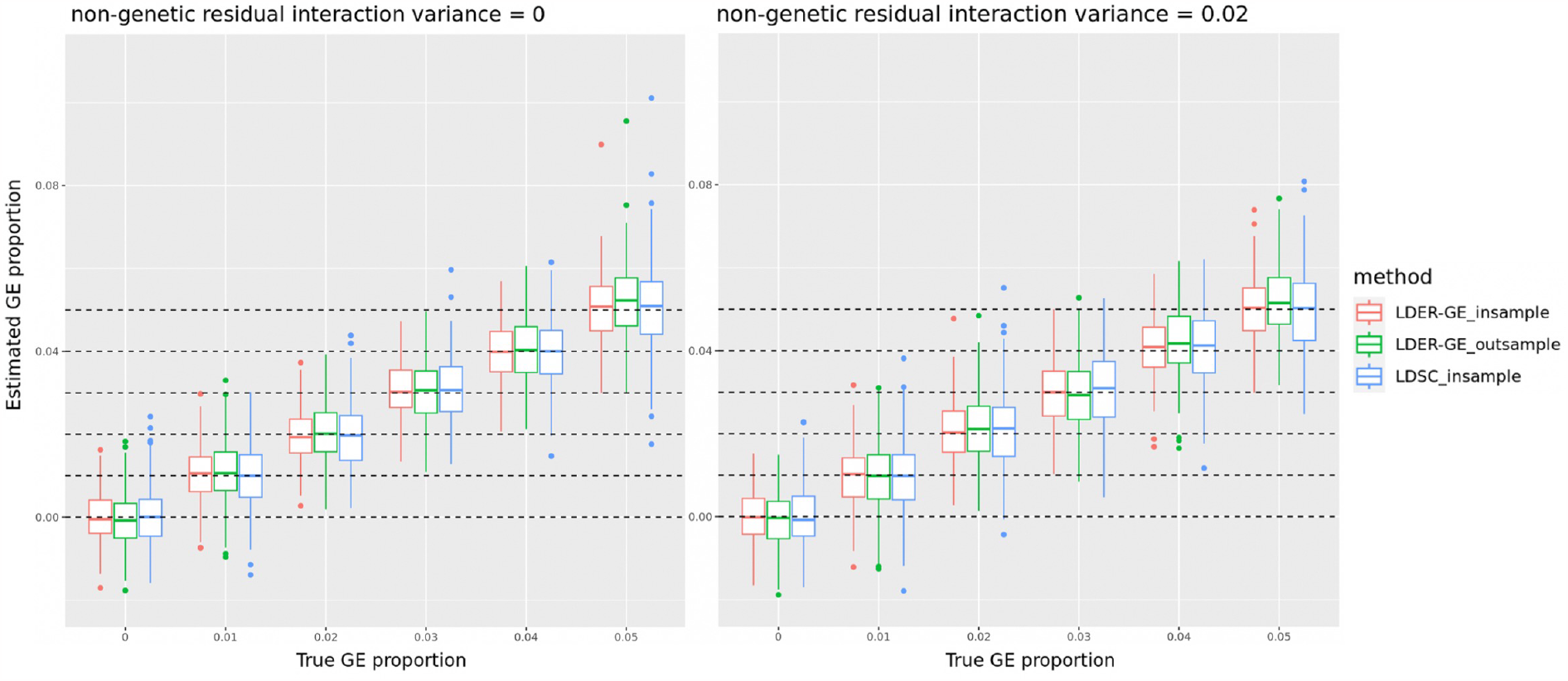
Histogram comparison of LDSC-based method and LDER-GE with in-sample and out-sample reference panel on simulations from real genotype panel, continuous phenotype.

The transformation from observed-scale variance to liability-scale variance, as achieved by the Roberston transformation^18^, relies on the normality assumption of the liability distribution. However, this assumption is violated with the non-normal GE term. Our simulations, considering various prevalence and GE interaction variance settings, suggest that the transformation provides reasonably unbiased results (S. Figure 1, Panels B, C, and D) when the disease is not rare, with a prevalence exceeding 10%, or when the GE interaction variance remains relatively small, at or below 5%. Nevertheless, in cases where the disease is rare, and the GE interaction variance is non-trivial (S. Figure 1, Panels A and B), the transformed liability-scale GE interaction variance estimate tends to be overestimated. This trend becomes more pronounced as the disease prevalence decreases. Despite the potential bias introduced by the Roberston transformation^18^ in the presence of the GE term, the type-I error rate for tests involving binary phenotypes remained well-controlled (Table 3) across varying prevalence settings. Our findings related to the estimation of GE liability-scale variance and the results of type-I error rate simulations for binary phenotypes align with a previous study^8^, except that LDER-GE achieved better estimation accuracy than LDSC-based method using in-sample and out-sample LD reference panels (S Table 1).

### Real data analysis using UKBB data

We examined 151 E-Y pairs involving 307,259 unrelated European ancestry individuals and a total of 966,766 genetic variants from the UK Biobank (UKBB). We employed both the LDER-GE and LDSC-based methods. Following Bonferroni correction, LDER-GE identified 35 significant pairs, of which 23 overlapped with the 25 pairs identified by the LDSC-based method (Figure 2). Further details about the 12 E-Y pairs exclusively identified by LDER-GE and the two E-Y pairs uniquely identified by the LDSC-based method can be found in S Table 2. Previous research has provided evidence of gene-age interaction effects on blood pressure through extensive GWAS data from three blood pressure consortia^19^ and linkage analysis^20^. In our analysis, LDER-GE successfully captured signals from both systolic blood pressure (SBP) and diastolic blood pressure (DBP), while the LDSC-based method failed to detect the DBP signal. Additionally, gene-sex interaction effects have been reported for traits such as height^21^, depression^22,23^ and cholesterol level^24^, all of which were exclusively detected by LDER-GE. In summary, the estimated values obtained using LDER-GE and the LDSC-based method exhibited strong overall consistency. However, we note that the standard error of LDER-GE was, on average, 21% lower than that of the LDSC-based method, a result consistent with our simulation findings. For a comprehensive overview of the analysis results for all 151 E-Y pairs, see S Table 3. Among the seven environmental covariates investigated, sex, Body Mass Index (BMI), and age exhibited relatively larger genome-level GE interaction effects and lower P-values compared to the other covariates (S Table 3). On the other hand, the pollution covariate pm2.5 did not exhibit statistically significant GE interactions across all tested phenotypes, including several lung-related traits. This can be a consequence of small GE interaction magnitudes and inaccuracy of air pollution indicator measurements.

**Figure 2:**
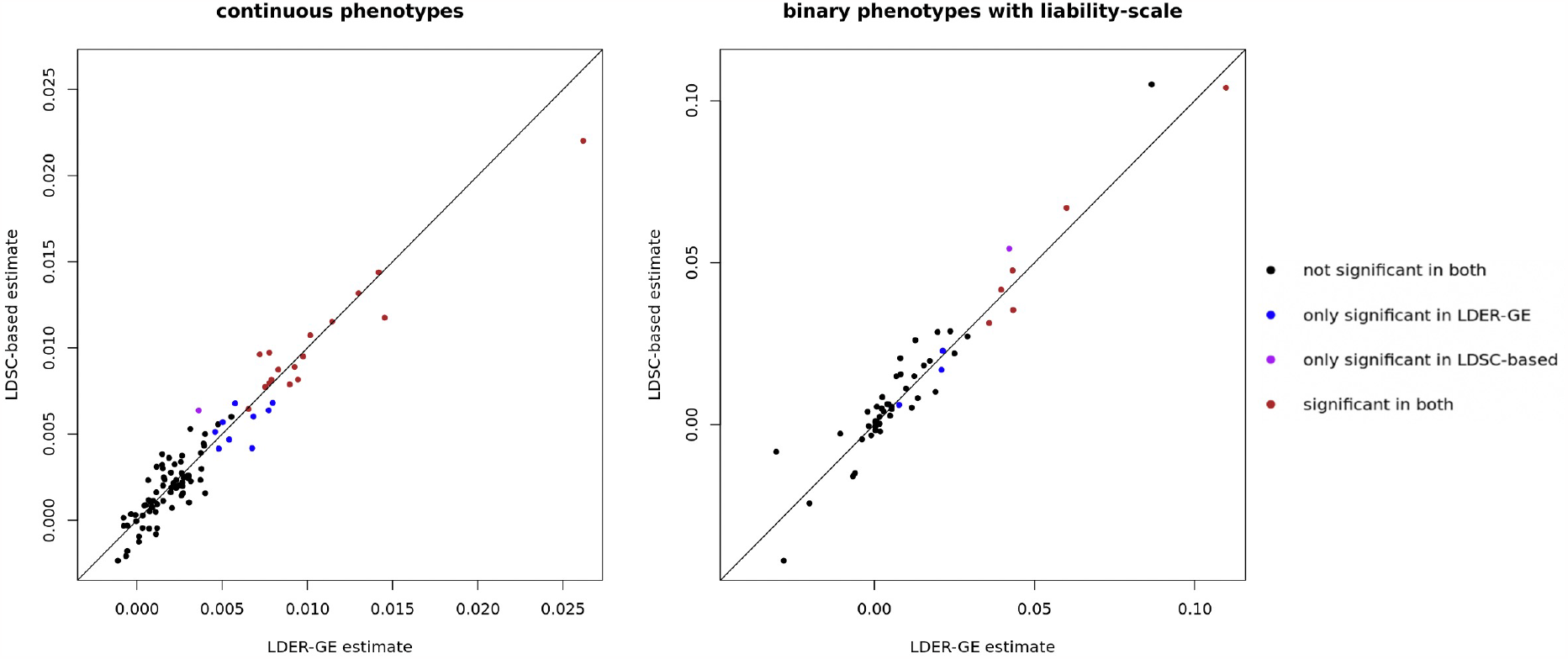
GE interaction variance estimates from LDSC-based method and LDER-GE of 151 environmental covariate-phenotype pairs in UKBB dataset. For binary phenotypes, GE interaction variance is reported on the liability-scale.

**Figure 3:**
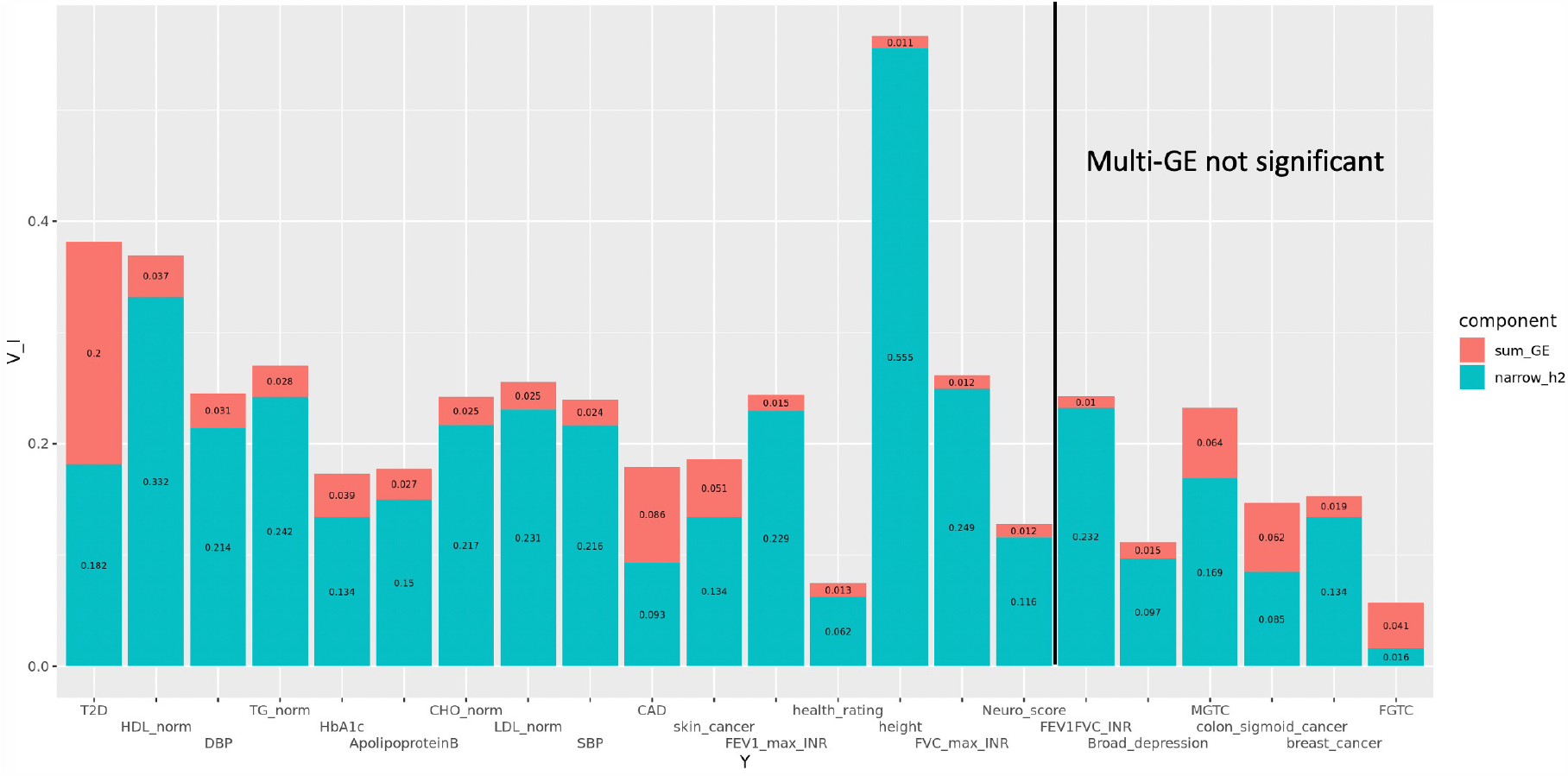
Phenotypic variance explained by narrow-sense heritability and aggregated multi-GE interactions for the 22 phenotypes, ordered by increasing P value of multi-GE interactions. Right of the solid black line, 6 phenotypes are not significant. For binary phenotypes, proportion is reported on the liability-scale. FGTC: Female genital tract cancer MGTC: Male genital tract cancer

For each of the 22 phenotypes, we estimated the aggregated multi-GE interaction variance using ordered covariates (age, sex, BMI, packed years of smoking, Townsend deprivation index, alcohol intake frequency). Post Bonferroni correction, LDSC-based method identified 13 phenotypes, while LDER-GE identified 16 phenotypes which covered all the 13 phenotypes discovered by LDSC-based method (S Table 4, S Figure 2). The additional 3 phenotypes identified are height, normalized FVC and neuroticism score, whose aggregated GE interaction variances were estimated around 1%. Notably, some phenotypes like depression and male genital tract cancer exhibited significant single-covariate GE interactions but were not detected with multi-GE interaction variance. This discrepancy may be due to noise introduced by other weak covariates during multi-covariate aggregation. Similar to single-covariate GE analysis, LDER-GE consistently yielded results comparable to the LDSC-based method in assessing multi-GE interaction variance but reported an average standard error 21% smaller than the LDSC-based method. We also estimated narrow-sense heritability h^2^ and compared it to the aggregated multi-GE interaction variance. Notably, 13 of the 22 phenotypes exhibited substantial multi-GE interaction variance, contributing more than 10% relatively to the narrow-sense heritability. For conditions like type-II diabetes (T2D) and coronary artery disease (CAD), the multi-GE interaction variance approached the magnitude of the narrow-sense heritability, providing valuable insights into disease etiology. However, we caution that when interpreting results for binary diseases, as the liability-scale transformation may introduce bias. We reversed the covariate order and used the set (alcohol intake frequency, Townsend deprivation index, packed years of smoking, BMI, sex, age) for the same analysis. It turned out that the order of covariates did not substantially affect the results (S Table 5).

## Discussion

In this study, we introduce LDER-GE to improve the precision of estimating GE interaction variance of complex traits using summary statistics. LDER-GE leverages full LD information from the LD panel while LDSC-based methods^7,8^ rely solely on the LD panel’s diagonal information. Our simulations and analysis of UK Biobank data demonstrate LDER-GE’s superiority over LDSC-based approaches in terms of estimation accuracy and root mean square error. LDER-GE’s improved accuracy enables the detection of more genome-level GE interactions that might go undetected by LDSC-based methods.

From real data analysis, sex and BMI had more detectable GE interaction effects over various health-related traits, consistent with results from other studies. For example, several studies reported that sex modifies genetic effects on lipid traits^25^, obesity^26,27^, and hypertension^28^ from different perspectives. BMI is known to be causal to multiple health-related traits such as T2D and hypertension^29,30^, part of which could be reasoned from the GE interaction effects^8^. While biological sex is almost fixed for most individuals in the population throughout the lifetime as well as its associated GE interaction variance, BMI varies during different life stages. The gene-BMI interaction study potentially brings additional significance to the clinical prevention or treatment to diseases such as T2D and hypertension, given their considerable GE interaction variance estimate. The higher statistical efficiency of LDER-GE allows us to better estimate the aggregated multi-GE interaction variance, which is comparable to narrow sense heritability, especially for T2D and Coronary Artery Disease (CAD). However, such interpretation must be accompanied with caution, due to the normality and additive effect assumption violations of Roberston transformation^18,31,32^. Empirically, simulations demonstrate the true aggregated multi-GE interaction variance is lower than the transformed liability-scale variance when the disease is rare (<=5%), but the magnitude difference is not considerable.

Our real data analysis highlighted the significant gene by SEX and gene by BMI interaction effects over various health-related traits, a finding consistent with diverse studies. For instance, sex has been shown to modify genetic effects on traits like lipid ^25^, obesity^26,27^, and hypertension^28^. Additionally, BMI, a known causal factor for multiple health-related traits such as T2D and hypertension^29,30^, may exert some of its influence through gene-environment (GE) interactions^8^. While biological sex remains relatively constant throughout an individual’s lifetime, BMI is subject to regulation. Investigating BMI’s GE interaction implications could have clinical significance, particularly in preventing or treating diseases like T2D and hypertension, given their substantial GE interaction variance estimates. LDER-GE’s enhanced statistical efficiency enables more accurate estimation of aggregated multi-GE interaction variance, which, notably, often approaches the magnitude of narrow-sense heritability, especially for T2D and CAD. However, it’s crucial to exercise caution in interpreting these findings, as they hinge on the normality and additive effect assumptions of the Roberston transformation^18,31,32^. Empirical simulations reveal that in cases of rare diseases (prevalence <= 5%), the true aggregated multi-GE interaction variance tends to be lower than the estimated liability-scale variance, though the difference in magnitude is not substantial.

As previously discussed^8^, it’s not recommended to directly estimate the non-genetic-residual-environment interaction variance from the intercept using the formula (intercept – 2*h_I_^2^)/2. This is because the intercept can be inflated by factors such as population stratification and other confounding effects, making it difficult to separate from the non-genetic-residual-environment interaction variance. When analyzing binary phenotypes, there’s the additional challenge of unknown prevalence differences between the sampling population and the target population, which can further inflate the estimated intercept^14^.

To expedite computation, we partitioned the entire genome into 1009 approximately independent blocks based on their LD relationships. We derived the block-wise LD matrix using a dataset of 307,259 individuals of European ancestry from the UK Biobank. Alternatively, we also calculated the out-sample LD matrix using 489 subjects from the 1000 Genomes project^33^, employing a linear shrinkage method^34,35^, and the 1703 genomic blocks from LDetect^36^. Typically, the 1000 Genomes project reference panel is utilized when in-sample LD information is unavailable. For readers’ convenience, both computed panels are accessible online. Our simulation results underscore the robustness of LDER-GE, whether LD panels are constructed from the UK Biobank or the 1000 Genomes project. However, it’s advisable to prioritize the UK Biobank reference when there’s a significant overlap between the variants in the GWIS input summary statistics and the UK Biobank reference panel, mainly due to its larger sample size.

As an extension of the LDSC-based method, LDER-GE inherits most of its limitations. Firstly, it assumes polygenic GE effects on the phenotype, and a violation of this assumption can result in underestimation of the variance component^37^. Secondly, our model does not differentiate between GE covariate correlations, potentially introducing estimation bias due to overadjustment^8^. Methods addressing such correlations are available^13^. Thirdly, LDER-GE has not been applied to case-control studies, which often involve oversampling of cases. While efforts are being made to address these limitations, future research could explore incorporating variant functional annotation or allele frequency information to enhance estimation^37^.

To summarize, LDER-GE utilizes full LD information and summary statistics to estimate the phenotypic variance explained by GE interactions more accurately than LDSC-based estimation methods. LDER-GE controls the computational burden and time well compared to methods that requires individual-level data input and can be employed to estimate multiple E-Y pairs of large sample size.

## Material & methods

### LDER-GE modelling & estimation

We consider the following model

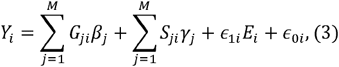

where *Y*_*i*_ is the phenotype for subject *i* already adjusted for fixed effects including the exposure covariate effects. *E*_*i*_ is the exposure covariate for subject *i*. Suppose there are *M* variants. *G*_*ji*_ is the *j*th variant for subject *i. S*_*ji*_ *= G*_*ji*_∗*E*_*i*_ is the GE interaction product term for variant *j* of subject *i*, □_*1i*_ is the non-genetic residual that has exposure interaction effect. □_0i_ is the residual independent from all other parts, β_*j*_ is the true additive genetic effect for variant *j*, γ_*j*_ is the true interaction effect for variant *j*. Following PEGION’s^7^ setting, we model β_*j*_ and γ_*j*_ using the following random effects model:

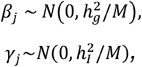

where 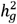 is the narrow-sense heritability and 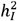 is the GE interaction variance that we are interested in estimating. And β_*j*_ and γ_*j*_ may or not be correlated. We model □_*0*_ and □_*1*_ using random effects model:

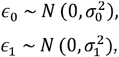

where 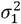 is the non-genetic environment interaction variance. Again, □_*0*_ and □_*1*_ may or may not be correlated.

Under the polygenic GE model, we derive (supplementary note 1), in matrix form, that

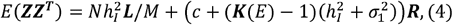

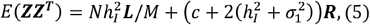

where ***Z*** is the GWIS Z-score vector, ***R*** is the LD matrix, ***L = R***^*T*^***R*** is the LD score matrix, *N* is the sample size of the GWAS summary statistics, *c* is the unconstrained intercept with potential inflation and ***K****(E)* is the kurtosis of the exposure covariate. In the case of *E* being standard normal, equation (4) reduces to the equation (5). Following the original LDER^15^ framework, we eigen-decompose the LD matrix as ***R*** *=* ***UDU***^*T*^, where ***U*** is the orthogonal matrix of eigenvectors and ***D*** is the diagonal eigen value matrix. Then we transform the original GWIS Z-score vector 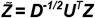 and have

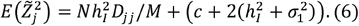

The transformed summary statistic vector contains all LD information, and the estimation efficiency is improved compared to LDSC-based methods as a consequence. The estimation task is accomplished using the iterative least squares and standard error is estimated using delete-block-wise jackknife.

To analyze binary outcomes, we transformed the observed-scale heritability to liability-scale heritability using Roberston transformation^18^. It has been pointed out that when the GE interaction variance is large, the normality assumption of the phenotype liability may be violated, resulting in biased results of Roberston transformation^8,38^. However, our simulation results showed that when the GE interaction variance proportion was small Roberston transformation still yielded reasonably accurate result (S Figure 1, S Table 1), consistent with previous studies^8^.

The regression weight of the transformed summary statistics vector takes the same form as LDER^15^ except the additional intercept inflation component (Supplementary note),

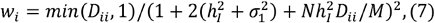

where *(1+ 2(h*^*2*^_*I*_ *+ σ*_*1*_^*2*^*) + Nh*^*2*^_*I*_*D*_*ii*_*/M)*^*2*^ is proportional to the variance of Z _*i*_ (supplementary note 2) and the shrinkage operation *min (D*_*ii*_,*1)* reduces the noise from big eigenvalues from LD matrix with lower sample sizes.

### UKBB data for simulation and real data analysis

The research conducted in this study utilized data from the UK Biobank Resource. The genomic partitioning and simulation analysis was conducted using UKBB dataset with application number 29900. The real data analysis was conducted using UKBB dataset with application number 32285. Detailed information regarding data access, ethical approval, quality control procedures, and phenotype definitions can be found in the supplementary note 2.

### Reference panel construction

We first took the intersection between UKBB^17^ imputed genotype panel, 1000 Genomes project^33^ genotype panel and hapmap3 project^39^ variant list, resulting in M = 396,330 common variants. Then, we partitioned the entire human genome into 1009 roughly independent blocks using the panel of 396,330 common variants from UKBB European ancestry (N=276,050). We partitioned the genome such that the linked SNP pairs (squared LD coefficient r^2^> 2/sqrt(276050) = 0.0038) are within 100 kilobases within each block. For simulations, in-sample reference panel was constructed using the intersected UKBB genotype panel (N=276,050, M = 396,330) and the 1009 genomic blocks. Out-sample reference panel was constructed using the same set of variants but from 1000 Genome project genotype panel (N=489, M = 396,330), with the genome partition being the 1703 genomic blocks generated previously^15^ for reducing the noise of low sample size. A linear shrinkage method^34,35^ was employed for out-sample reference panel construction to further reduce the noise. For UKBB real data analysis, the in-sample reference panel was constructed using the union of UKBB imputed genotype panel and UKBB array genotype panel, intersected with hapmap3 project^39^ variant list (N=307,259, M = 966,766) and the 1009 genomic blocks. For real data analysis, the variant inclusion criteria was (a): imputation score > 0.8; (b): minor allele frequency > 0.05; (c): missing rate < 0.01; (d): Hardy-Weinberg Equilibrium P-value > 5*10^-8^. The details of quality control procedure of simulation dataset and real analysis dataset can be found in the supplementary materials.

### Simulations

The data generation process followed equation X, with narrow-sense heritability h^2^_g_ fixed at 0.2, GE interaction contribution proportion h^2^_I_ varying from 0 to 0.05, non-genetic-residual-covariate interaction variance σ_1_^2^ being 0 or 0.02, E is standard normal:

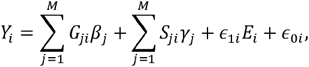

For each simulation from the pool of intersected UKBB genotype panel (N=276,050, M = 396,330), we randomly chose 50,000 subjects and 19,816 (5%) causal GE variants for data generation, association analysis (linear regression on M = 396,330 variants using PLINK2^40^) and GE interaction variance estimation analysis. Each parameter combination had 300 replicated simulations. To simulate the binary outcome, we used the liability model based on corresponding critical cutoff with respect to the specified population prevalence.

### Single-covariate GE interaction variance analysis of UKBB

We ran GWIS analysis through “--glm interaction --variance-standardize” command of PLINK2^40^ pre-adjusted for age, sex, 40 genetic PCs and the specific environmental covariate of interest if not age or sex using linear regression. We analyzed 22 phenotypes and 7 environmental covariates, resulting in a total of 151 (154 - 3) E-Y pairs with the 3 sex-specific phenotypes. The 22 phenotypes included 14 continuous phenotypes and 8 binary phenotypes. LDER-GE and LDSC-based analysis were conducted using the resulted GWIS summary statistics and pre-computed LD information.

### Aggregated multi-covariate GE interaction variance analysis of UKBB

Suppose there is a covariate set of interest (A, B, C, …), we first run linear regression of B∼A to get residuals of B net A: B|A, being independent from A, and we run another linear regression of C∼A+B to get residuals of C net A and B: C|A & B, being independent from A and B. We continue the process until all residuals are independent from each other. We then run single-covariate GE interaction variance analysis on each residuals the same way but preadjust for age, sex, 40 genetic PCs and all covariates in the set excluding age and sex. By eliminating the dependency of covariates, the estimated single-covariate GE interaction variances are independent, and we summed up the estimated GE interaction variances and their variances of estimation to conduct straightforward statistical test. We explored the set (age, sex, BMI, packed years of smoking, Townsend deprivation index, Alcohol intake frequency) to capture more missing heritability explained by GE interactions because these 6 covariates yielded nonminimally significant GE interaction signals at P<0.05 on more than three phenotype (S table 6). The narrow-sense heritability of each phenotype was estimated using LDER and the main genetic effect GWAS summary statistics of the same UKBB cohort as GE analysis.

## Abbreviations

GE: gene-environment
GWIS: genome-wide interaction scan
LD: linkage disequilibrium
SNP: single nucleotide polymorphism
E-Y pair: environmental covariate phenotype pair

