## Supplementary material for "LDER-GE estimates phenotypic variance component of gene-environment interactions in human complex traits accurately with GE interaction summary statistics and full LD information": table 1-4

| H_I_^2 a^ | method | mean | Precision^b^ | RMSE^c^ |
| --- | --- | --- | --- | --- |
| ***0*** | ***LDER-GE_insample*** | ***0.000*** | ***168.40*** | ***0.0059*** |
| 0 | LDER-GE_outsample | -0.001 | 152.58 | 0.0066 |
| 0 | LDSC_insample | 0.000 | 139.80 | 0.0072 |
| ***0.01*** | ***LDER-GE_insample*** | ***0.010*** | ***152.48*** | ***0.0066*** |
| 0.01 | LDER-GE_outsample | 0.010 | 134.28 | 0.0074 |
| 0.01 | LDSC_insample | 0.010 | 126.84 | 0.0079 |
| ***0.02*** | ***LDER-GE_insample*** | ***0.020*** | ***151.09*** | ***0.0066*** |
| 0.02 | LDER-GE_outsample | 0.021 | 136.29 | 0.0074 |
| 0.02 | LDSC_insample | 0.020 | 121.81 | 0.0082 |
| ***0.03*** | ***LDER-GE_insample*** | ***0.030*** | ***140.87*** | ***0.0071*** |
| 0.03 | LDER-GE_outsample | 0.030 | 128.21 | 0.0078 |
| 0.03 | LDSC_insample | 0.031 | 116.09 | 0.0086 |
| ***0.04*** | ***LDER-GE_insample*** | ***0.040*** | ***147.31*** | ***0.0068*** |
| 0.04 | LDER-GE_outsample | 0.041 | 126.79 | 0.0080 |
| 0.04 | LDSC_insample | 0.040 | 119.18 | 0.0084 |
| ***0.05*** | ***LDER-GE_insample*** | ***0.050*** | ***128.75*** | ***0.0078*** |
| 0.05 | LDER-GE_outsample | 0.052 | 115.67 | 0.0089 |
| 0.05 | LDSC_insample | 0.050 | 102.15 | 0.0098 |

Table 1: Simulation efficiency of LDER-GE and LDSC-based methods for estimating continuous phenotype GE interaction variance proportion. The summarized results for different non-genetic residual-environment interaction variance (0 or 0.02) are merged. Each row has 2*300 = 600 replications.

H_I_^2 a^: True GE interaction variance proportion.

Precision^b^: 1/Empirical standard deviation.

RMSE^c^: Root mean squared error rate.

***Highlighted method:*** highest precision and lowest RMSE among the three methods.

| Methods | σ_1_^2 a^ | Type-I error Rate |
| --- | --- | --- |
| LDER-GE_insample | 0 | 0.0500 |
|  | 0.02 | 0.0480 |
|  | **Average** | **0.0490** |
| LDER-GE_outsample | 0 | 0.0600 |
|  | 0.02 | 0.0500 |
|  | **Average** | **0.0550** |
| LDSC_insample | 0 | 0.0633 |
|  | 0.02 | 0.0400 |
|  | **Average** | **0.0517** |

Table 2: Type-I error Rate at 0.05 level for LDER-GE and LDSC-based method for continuous phenotype simulation scenarios, each scenario with 300 replications.

σ_1_^2 a^: non-genetic residual-environment interaction variance.

| method | Prevalence | | Type-I error Rate |
| --- | --- | --- | --- |
| LDER-GE_insample | 0.05 | 0.0400 | |
|  | 0.1 | 0.0533 | |
|  | 0.2 | 0.0500 | |
|  | 0.3 | 0.0567 | |
|  | **Average** | **0.0500** | |
| LDER-GE_outsample | 0.05 | 0.0467 | |
|  | 0.1 | 0.0600 | |
|  | 0.2 | 0.0533 | |
|  | 0.3 | 0.0567 | |
|  | **Average** | **0.0541** | |
| LDSC_insample | 0.05 | 0.0533 | |
|  | 0.1 | 0.0533 | |
|  | 0.2 | 0.0467 | |
|  | 0.3 | 0.0533 | |
|  | **Average** | **0.0517** | |

Table 3: Type-I error Rate at 0.05 level for LDER-GE and LDSC-based method for binary phenotype simulation scenarios, each scenario with 300 replications.

| Y | GE_variance | | SE | P | h^2^ | | GE_variance/h^2^ |
| --- | --- | --- | --- | --- | --- | --- | --- |
| T2D | 2.00E-01 | 1.95E-02 | | 1.33E-24 | 1.82E-01 | 1.10E+00 | |
| HDL_norm | 3.72E-02 | 3.81E-03 | | 1.60E-22 | 3.32E-01 | 1.12E-01 | |
| DBP | 3.11E-02 | 3.32E-03 | | 7.32E-21 | 2.14E-01 | 1.45E-01 | |
| TG_norm | 2.84E-02 | 3.43E-03 | | 1.34E-16 | 2.42E-01 | 1.17E-01 | |
| HbA1c | 3.90E-02 | 4.78E-03 | | 3.74E-16 | 1.34E-01 | 2.91E-01 | |
| ApolipoproteinB | 2.75E-02 | 3.40E-03 | | 5.83E-16 | 1.50E-01 | 1.83E-01 | |
| CHO_norm | 2.50E-02 | 3.34E-03 | | 6.93E-14 | 2.17E-01 | 1.15E-01 | |
| LDL_norm | 2.52E-02 | 3.38E-03 | | 9.70E-14 | 2.31E-01 | 1.09E-01 | |
| SBP | 2.38E-02 | 3.26E-03 | | 2.70E-13 | 2.16E-01 | 1.10E-01 | |
| CAD | 8.59E-02 | 1.89E-02 | | 5.19E-06 | 9.33E-02 | 9.21E-01 | |
| skin_cancer | 5.11E-02 | 1.18E-02 | | 1.49E-05 | 1.34E-01 | 3.80E-01 | |
| FEV1_max_INR | 1.46E-02 | 3.69E-03 | | 7.44E-05 | 2.29E-01 | 6.36E-02 | |
| health_rating | 1.29E-02 | 3.28E-03 | | 8.94E-05 | 6.21E-02 | 2.07E-01 | |
| height | 1.10E-02 | 3.09E-03 | | 3.80E-04 | 5.55E-01 | 1.98E-02 | |
| FVC_max_INR | 1.21E-02 | 3.47E-03 | | 5.09E-04 | 2.49E-01 | 4.84E-02 | |
| Neuro_score | 1.18E-02 | 3.66E-03 | | 1.29E-03 | 1.16E-01 | 1.02E-01 | |
| FEV1FVC_INR | 1.00E-02 | 3.40E-03 | | 3.14E-03 | 2.32E-01 | NA | |
| Broad_depression | 1.45E-02 | 4.96E-03 | | 3.45E-03 | 9.69E-02 | NA | |
| male_genital_tract_cancer | 6.37E-02 | 2.30E-02 | | 5.63E-03 | 1.69E-01 | NA | |
| colon_sigmoid_cancer | 6.22E-02 | 3.49E-02 | | 7.45E-02 | 8.49E-02 | NA | |
| breast_cancer | 1.89E-02 | 1.59E-02 | | 2.35E-01 | 1.34E-01 | NA | |
| female_genital_tract_cancer | 4.11E-02 | 5.25E-02 | | 4.34E-01 | 1.61E-02 | NA | |

Table 4: aggregated multi-GE interaction variance and narrow sense heritability (h^2^) comparison of the 22 phenotypes.
