## Supplemental file 1 for "LDER-GE estimates phenotypic variance component of gene-environment interactions in human complex traits accurately with GE interaction summary statistics and full LD information"

### 1 Main derivation

We set up the model as [1].

$$Y_i = \sum_{j=1}^{M} G_{ji}\beta_j + \sum_{j=1}^{M} S_{ji}\gamma_j + \epsilon_{1i}E_i + \epsilon_{0i} \tag{1}$$

Where $Y_i$ is the phenotype for subject i already adjusted for fixed effect including for the exposure covariate. $E_i$ is the exposure covariate for subject i. Suppose there are M variant. $G_{ji}$ is the jth variant for subject i. $S_{ji} = G_{ji} * E_i$ is the GE interaction product term for variant j of subject i. $\epsilon_{1i}$ is the non-genetic residual that has exposure interaction effect. $\epsilon_{0i}$ is the residual independent from all other parts.$\beta_j$ is the true additive effect for variant j. $\gamma_j$ is the true interaction effect for variant j.

In matrix form, it's written as

$$\boldsymbol{Y} = \boldsymbol{G\beta} + \boldsymbol{S\gamma} + \boldsymbol{\epsilon_1 E} + \boldsymbol{\epsilon_0} \tag{2}$$

We model $\beta_j$ and $\gamma_j$ using random effect model:

$$\begin{pmatrix}\beta_j \\ \gamma_j\end{pmatrix} \sim N\left[\begin{pmatrix}0 \\ 0\end{pmatrix}, \begin{pmatrix}h_g^2/M & \rho_{gI}/M \\ \rho_{gI}/M & h_I^2/M\end{pmatrix}\right] \tag{3}$$

Where $h_g^2$ is the narrow-sense heritability and $h_I^2$ is the GE interaction contribution proportion that we are interested at. $\rho_{gI}$ models the genome-level covariance between the additive effect and GE interaction effect.
We model $\epsilon_0$ and $\epsilon_1$ using random effect model:

$$\begin{pmatrix}\epsilon_{0_i} \\ \epsilon_{1_i}\end{pmatrix} \sim N\left[\begin{pmatrix}0 \\ 0\end{pmatrix}, \begin{pmatrix}\sigma_0^2 & \rho_{0,1} \\ \rho_{0,1} & \sigma_1^2\end{pmatrix}\right] \tag{4}$$

We assume that the genetic part and residual part are independent.

Suppose all variants and the exposure variable are standardized.
Then the interaction effect estimate for variant j is:

$$\begin{aligned}\hat{\gamma_j} &= (\boldsymbol{S_{j.}^T S_{j.}})^{-1}\boldsymbol{S_{j.}^T Y} \\ &= \boldsymbol{S_{j.}^T Y}/n\end{aligned} \tag{5}$$

The first equality holds because after standardization $\text{var}(S_{j.}) = 1$
And the Z score of the interaction effect estimate for variant j is:

$$Z_{jI} = \boldsymbol{S_{j.}^T Y}/\sqrt{n} \tag{6}$$

Then for two variant j and k, the expected interaction effect estimate Z score product is:

$$\begin{aligned}\mathbb{E}(Z_{jI}Z_{kI}) &= \mathbb{E}(\mathbb{E}(Z_{jI}Z_{kI}|\boldsymbol{\beta}, \boldsymbol{\gamma}, \boldsymbol{\epsilon_0}, \boldsymbol{\epsilon_1})) \\ &= \mathbb{E}(\mathbb{E}(\boldsymbol{S_{j.}^T Y S_{k.}^T Y}/n|\boldsymbol{\beta}, \boldsymbol{\gamma}, \boldsymbol{\epsilon_0}, \boldsymbol{\epsilon_1})) \\ &= \mathbb{E}(\boldsymbol{S_{j.}^T(G\beta + S\gamma + \epsilon_1 E + \epsilon_0)S_{k.}^T(G\beta + S\gamma + \epsilon_1 E + \epsilon_0)})/n \\ &= \mathbb{E}(\boldsymbol{S_{j.}^T(G\beta)S_{k.}^T(G\beta)})/n + \mathbb{E}(\boldsymbol{S_{j.}^T(S\gamma)S_{k.}^T(S\gamma)})/n \\ &+ \mathbb{E}(\boldsymbol{S_{j.}^T(\epsilon_1 E)S_{k.}^T(\epsilon_1 E)})/n + \mathbb{E}(\boldsymbol{S_{j.}^T(\epsilon_0)S_{k.}^T(\epsilon_0)})/n \\ &= h_g^2 M r_{jk}/M + h_I^2/M * ((n^2 + 2nK(E) - 2n)\ell_{jk} + nMK(E)r_{jk})/n \\ &+ \sigma_1^2 K(E) r_{jk} + \sigma_0^2 r_{jk} \\ &\approx h_I^2 n\ell_{jk}/M + h_g^2 r_{jk} + h_I^2 K(E) r_{jk} + \sigma_1^2 K(E) r_{jk} + \sigma_0^2 r_{jk} \\ &= h_I^2 n\ell_{jk}/M + (1 + (K(E) - 1)(h_I^2 + \sigma_1^2))r_{jk}\end{aligned} \tag{7}$$

where $r_{jk}$ is the true correlation between the SNP j and k, and $\ell_{jk} = \sum_{t=1}^{M} r_{jt}r_{kt}$ is the off diagonal element of the LD score matrix. $K(E)$ is the forth moment of exposure E. The details of the fifth equality are in supplementary.

In matrix form, the above quantity can be written as

$$\mathbb{E}(\boldsymbol{ZZ^T}) = Nh_I^2\boldsymbol{L}/M + (c + 2(h_I^2 + \sigma_1^2))\boldsymbol{R} \tag{8}$$

where $\boldsymbol{L} = \boldsymbol{R^T R}$ is the LD score matrix and c is the unconstrained intercept.

### 2 Details

This section provides the derivation details.
For equation (7) equality 5 there are 4 parts:

$$\begin{aligned} &\mathbb{E}(\boldsymbol{S_{j.}^T}\boldsymbol{(G\beta)}\boldsymbol{S_{k.}^T}\boldsymbol{(G\beta)}/n + \mathbb{E}(\boldsymbol{S_{j.}^T}\boldsymbol{(S\gamma)}\boldsymbol{S_{k.}^T}\boldsymbol{(S\gamma)})/n \\ &+\mathbb{E}(\boldsymbol{S_{j.}^T}\boldsymbol{(\epsilon_1 E)}\boldsymbol{S_{k.}^T}\boldsymbol{(\epsilon_1 E)})/n + \mathbb{E}(\boldsymbol{S_{j.}^T}\boldsymbol{(\epsilon_0)}\boldsymbol{S_{k.}^T}\boldsymbol{(\epsilon_0)})/n \\ &= h_g^2 M r_{jk}/M + h_I^2/M * ((n^2+2nK(E)-2n)\ell_{jk} + nMK(E)r_{jk})/n \\ &+ \sigma_1^2 K(E) r_{jk} + \sigma_0^2 r_{jk} \end{aligned} \tag{9}$$

The first part:

$$\begin{aligned} &\mathbb{E}(\boldsymbol{S_{j.}^T}\boldsymbol{(G\beta)}\boldsymbol{S_{k.}^T}\boldsymbol{(G\beta)})/n \\ &= \mathbb{E}(\boldsymbol{S_{j.}^T}\boldsymbol{(G\beta)}\boldsymbol{(G\beta)^T}\boldsymbol{S_{k.}})/n \\ &= h_g^2 M/M * \boldsymbol{E}(\boldsymbol{S_{j.}^T}\boldsymbol{S_{k.}})/n \\ &= h_g^2 M/M * \boldsymbol{E}(\boldsymbol{G_{j.}^T}\boldsymbol{G_{k.}})/n \\ &= h_g^2 r_{jk} \end{aligned} \tag{10}$$

To start the derivation for the second part, we need to review some derivations from the paper [2].

$$\boldsymbol{G_{j.}^T}\boldsymbol{G}\boldsymbol{G^T}\boldsymbol{G_{k.}} = n^2 \sum_{t=1}^{M} \tilde{r_{jt}}\tilde{r_{kt}} \tag{11}$$

$$\mathbb{E}(\sum_{t=1}^{M} \tilde{r_{jt}}\tilde{r_{kt}}) \approx \ell_{jk} + M r_{jk}/n \tag{12}$$

Then, for our target quantity

$$\begin{aligned}
&\mathbb{E}(\boldsymbol{S_{j.}^T(S\gamma)S_{k.}^T(S\gamma)})/n \\
&= h_I^2/M * \boldsymbol{E}(\boldsymbol{S_{j.}^T S S^T S_{k.}})/n \\
&= h_I^2/M * \boldsymbol{E}(\sum_{t=1}^{M} \boldsymbol{S_{j.}^T S_{t.} S_{k.}^T S_{t.}})/n \\
&= h_I^2/M * \boldsymbol{E}(\sum_{t=1}^{M}(\sum_{i=1}^{N} G_{ji}G_{ki}E_i^2) * (\sum_{i=1}^{N} G_{ti}G_{ki}E_i^2))/n \\
&= h_I^2/M * \boldsymbol{E}(\sum_{t=1}^{M}(\sum_{i=1}^{N}(G_{ji}G_{ki}G_{ti}^2E_i^4) + 2 * \sum_{i\neq w}(G_{ji}G_{ti}G_{jw}G_{tw}E_i^2E_w^2)))/n \\
&= h_I^2/M * (K(E)-1)\boldsymbol{E}(\sum_{t=1}^{M}(\sum_{i=1}^{N}(G_{ji}G_{ki}G_{ti}^2))/n + h_I^2/M * \boldsymbol{E}(\boldsymbol{G_{j.}^T G G^T G_{k.}})/n \\
&\approx h_I^2/M * [(K(E)-1)(2n\ell_{jk} + nMr_{jk})/n + n^2(\ell_{jk} + Mr_{jk}/n)/n] \\
&= h_g^2 M r_{jk}/M + h_I^2/M * ((n^2 + 2nK(E) - 2n)\ell_{jk} + nMK(E)r_{jk})/n
\end{aligned} \tag{13}$$

The derivation of the sixth equality can be found in equation 16 and 17.
The third part:

$$\begin{aligned}
&\mathbb{E}(\boldsymbol{S_{j.}^T(\epsilon_1 E)S_{k.}^T(\epsilon_1 E)})/n \\
&= \sigma_1^2\mathbb{E}(\boldsymbol{S_{j.}^T E E^T S_{k.}})/n \\
&= \sigma_1^2 K(E) r_{jk}
\end{aligned} \tag{14}$$

The forth part:

$$\begin{aligned}
&\mathbb{E}(\boldsymbol{S_{j.}^T(\epsilon_0)S_{k.}^T(\epsilon_0)})/n \\
&= \sigma_0^2\boldsymbol{E}(\boldsymbol{S_{j.}^T S_{k.}})/n \\
&= \sigma_0^2 r_{jk}
\end{aligned} \tag{15}$$

To show the sixth equality in equation 13, we model three standardized variants $G_{ji}, G_{ki}, G_{ti}$ for one subject i as MVN distribution.

$$\begin{pmatrix} G_{ji} \\ G_{ki} \\ G_{ti} \end{pmatrix} \sim N \left[ \begin{pmatrix} 0 \\ 0 \\ 0 \end{pmatrix}, \quad \begin{pmatrix} 1 & r_{jk} & r_{jt} \\ r_{jk} & 1 & r_{kt} \\ r_{jt} & r_{kt} & 1 \end{pmatrix} \right] \tag{16}$$

Then, by low of total expectation,

$$\begin{aligned}
&\mathbb{E}(G_{ji}G_{ki}G_{ti}^2) \\
&= \mathbb{E}(\boldsymbol{E}(G_{ji}G_{ki}G_{ti}^2|G_{ti}) \\
&= 2r_{jt}r_{kt} + r_{jk}
\end{aligned} \tag{17}$$
