## Supplemental file 2 for "LDER-GE estimates phenotypic variance component of gene-environment interactions in human complex traits accurately with GE interaction summary statistics and full LD information"

### Supplementary note two of LDER-GE

#### Real data analysis UKBB dataset quality control and phenotype definition

Data access and ethical approval. This research was conducted using the UK Biobank Resource (application number 32285). The UK Biobank has approval from the Northwest Multi-centre Research Ethics Committee (MREC) to obtain and disseminate data and samples from the participants, and these ethical regulations cover the work in this study. Written informed consent was obtained from all participants before enrollment in the study, which was conducted in accordance with the principles of the Declaration of Helsinki. Details can be found at www.ukbiobank.ac.uk/ethics. The present analyses were approved by the Human Investigations Committee at Yale University (2000026836).

UK Biobank data. Participants were enrolled in the UK Biobank, and data accessed under an approved agreement (application ID 32285). Genotypes were assayed on either the UK BiLEVE array or the UK Biobank Axiom array with 733,332 autosomal variants overlapping between the two arrays. Genotype imputation based on Haplotype Reference Consortium and 1000 Genome project panel yielded 93 million variants for each subject. Subject and variant quality control is described in Supplemental Table S7, with subjects limited to those who self-defined and were genetically confirmed as White British (field 22006) and with a call rate >99% variants. We limited the dataset by using the indicator variable (field 22021) which indicates that the subjects were unrelated (at least 3$^{rd}$ degree relatives) and included in the original principal components (PC) calculation. This resulted in a subset of 307,259 unrelated individuals for analysis. Final variant quality control steps were call rate >99%, Hardy-Weinberg Equilibrium (HWE) p-value > $5 \times 10^{-8}$, and minor allele frequency (MAF) > 0.05.

Phenotype and environmental covariate definition

Asthma: ICD-10 code (field 41270, J45 or J46) or self-reported diagnosis by a doctor (field 6152). Individuals with autoimmune conditions were excluded from the controls (field 20002, self-reported diagnosis of an autoimmune disease[1], or self-report of sarcoidosis diagnosis by doctor in field 22133). Individuals who did not report a diagnosis of asthma at the recruitment visit but did report a diagnosis at a later visit were also excluded from both the cases and controls. There were 47,719 cases and 313,086 controls.

BMI: (field 21001) excluded women pregnant at time of recruitment (field 3140). BMI was transformed by rank-based inverse normal transformation in R.

Systolic blood pressure: measured by taking two automated (field 4080; N=472,254 subjects) or manual (field 94; N=43,795 subjects) readings. When there were automated and manual readings available, the automated readings were used. The average of the two readings was used as the SBP value. To account for the use of blood pressure lowering medications, we added 15mmHg to the SBP value for all subjects taking one or more blood pressure lower medications[2]. SBP was transformed by rank-based inverse normal transformation in R.

Diastolic blood pressure: measured by taking two automated (field 4080; N=472,254 subjects) or manual (field 94; N=43,795 subjects) readings. When there were automated and manual readings available, the automated readings were used. The average of the two readings was used as the DBP value. To account for the use of blood pressure lowering medications, we added 10mmHg to the DBP value for all subjects taking one or more blood pressure lower medications[2]. SBP was transformed by rank-based inverse normal transformation in R.

ApolipoproteinB: filed 30640.

Glucose: field 30740.

HbA1c: field 30750.

Health_rating: Overall health rating, field 2178. We transformed the categorical scale to integer scale: “Excellent” -> 4; “Good” -> 3; “Fair” -> 2; “Poor” -> 1. “Do not know” and “Prefer not to answer” were excluded.

Height: standing height, field 50.

Neuro_score: Neuroticism score, field 20127.

CAD: Coronary artery disease, ICD-10 code (field 41270, I210-I214, I219, K401-K404, K411-K414, K451-K455, K491, K492, K498, K499, K502, K751-K754, K758, K759).

T2D: type II diabetes, ICD-9 code of K51; ICD-10 code of E11; or self-reported diagnosis by a doctor at ≥ 30 years of age (fields 2,443 and 2,976). Individuals with type 1 diabetes [self-reported diabetes that occurred <30 years of age or E10] or gestational diabetes [self-report (field 4,011) or O24] were excluded from both cases and controls.

HDL_norm: HDL cholesterol levels were obtained from field 30760 and inverse rank normalized.

LDL_norm: field 30780. LDL cholesterol was adjusted for those individuals who reported taking one of five cholesterol lowering drugs (field 20003; Simvastatin, atorvastatin, rosuvastatin, pravastatin, Fluvastatin) by dividing the measured LDL value by 0.7[3]. The resulting value was the inverse rank normalized.

TG_norm: Triglyceride levels were obtained from field 30870 and inverse rank normalized.

CHO_norm: field 30690. Totalholesterol was adjusted for those individuals who reported taking one of five cholesterol lowering drugs (field 20003; Simvastatin, atorvastatin, rosuvastatin, pravastatin, Fluvastatin) by dividing the measured value by 0.8[3]. The resulting value was the inverse rank normalized.

Breast_caner: Cancer code field 20001 to be 1002 (breast) or ICD-10 code (field 41270, C50).

colon_sigmoid_cancer: Cancer code field 20001 to be 1022 (colon cancer/sigmoid cancer) or ICD-10 code (field 41270, C18).

female_genital_tract_cancer: Cancer code field 20001 to be 1037 (female genital tract cancer) or ICD-10 code (field 41270, C51-C58).

male_genital_tract_cancer: Cancer code field 20001 to be 1038 (male genital tract cancer) or ICD-10 code (field 41270, C60-C63).

skin_cancer: Cancer code field 20001 to be 1003 (skin_cancer) or ICD-10 code (field 41270, C43-C44).

FEV1_max_INR: Forced expiratory volume in 1-second, data field 3063. Took maximum FEV1 reads of each subject and normalized them to Z-scores.

FVC_max_INR: Forced vital capacity, data field 3062. Took maximum FVC reads of each subject and normalized them to Z-scores.

FEV1FVC_INR: Took (maximum FEV1 reads / maximum FVC reads) and normalized them to Z-scores.

Broad_depression: broadly defined depression definition[4]. Seen doctor (GP) for nerves, anxiety, tension or depression, data field 2090. Or seen a psychiatrist for nerves, anxiety, tension or depression, data field 2100. Or ICD-10 code (field 41270, F32-34, F38-F39).

AGE: Age at recruitment, field 21022.

Alcohol_inake_frequency: Alcohol inake frequency, field 1558. We transformed the categorical scale to integer scale: “Daily or almost daily” -> 1; “Three or four times a week” -> 2; “Once or twice a week” -> 3; “One to three times a month” -> 4. “Special occasions only” -> 5; “Never” -> 6; “Prefer not to answer” was excluded. As in https://biobank.ndph.ox.ac.uk/ukb/coding.cgi?id=100402.

NO2: Nitrogen dioxide air pollution 2010, field 24003.

Pm2.5: Particulate matter 2.5 air pollution 2010, field 24006.

SEX: field 31.

smoking_years: packed years of smoking, field 20161. Every subject with missing values were set to be 0.

townsendscore: Townsend deprivation index at recruitment, field 22189.

**Supplementary table 7: initial array genotyping and subject quality control**

| | # variants | # variants removed in this step | # subjects | # subjects removed in this step |
|---|---|---|---|---|
| Step 1: Initial variant QC | | | | |
| Genotyped variants | 805,426 | | | |
| Autosomal variants | 784,256 | 21,170 | | |
| Covered by both arrays | 733,322 | 50,934 | | |
| Batch level qc | 687,004 | 46,318 | | |
| SNPs only (indels removed) | 674,489 | 12,515 | | |
| Step 2: Subject QC[1] | | | | |
| Genotypes available | | | 488,377 | |
| Phenotypes available | | | 488,282 | 95 |
| Genetic and reported sex match | | | 487,910 | 372 |
| Sex chromosomes non-XX XY | | | 487,440 | 470 |
| Outliers in heterozygosity/missing rate | | | 486,477 | 963 |
| "Caucasian" (f.22006) | | | 408,186 | 78,291 |
| Individual call rate > 99% | | | 366,752 | 41,434 |
| Unrelated | | | 307,259 | 59,493 |

[1]Subject QC was performed using the 674,489 variants

#### Simulation UKBB dataset quality control

The same set of N=276,050 subjects in the previous study[5] were used here: UKBB subjects with self-reported European ancestry (f.22006) and not related to any other subjects in the UKBB dataset (f.22021). We took the intersection of the UKBB imputed 93 million variants, hapmap3 list variants and variants in the 1000 Genome project, with the following quality control procedure: imputation score > 0.3, call rate >99%, Hardy-Weinberg Equilibrium (HWE) p-value > $5\times10^{-4}$, and minor allele frequency (MAF) > 0.05. This resulted in the set of 396,330 variants.

#### Derivation of regression weights

Following the original model, variant-level GE effect size $\gamma_j \sim iid\ N(0, h_I^2/M)$ across M variants, the LD matrix eigen decomposition $\mathbf{R} = \mathbf{U}\mathbf{D}\mathbf{U}^T$, and the transformation of Z vectors $\tilde{\mathbf{Z}} = \mathbf{D}^{-1/2}\mathbf{U}^T\mathbf{Z}$, we derive that the conditional distribution of transformed Z scores are $\tilde{Z} \mid \boldsymbol{\gamma} \sim N(\sqrt{N}D^{1/2}U^T\boldsymbol{\gamma}, (\mathbf{1} + 2(h_I^2 + \sigma_1^2))\boldsymbol{I})$.

The expectation and covariance of the marginal distribution of transformed Z scores are

$E(\tilde{Z}) = E\left(E(\tilde{Z}|\boldsymbol{\gamma})\right) = \mathbf{0}$ and $Cov(\tilde{Z}) = E(Cov(\tilde{Z} \mid \boldsymbol{\gamma})) + Cov(E(\tilde{Z} \mid \boldsymbol{\gamma})) = (\mathbf{1} + 2(h_I^2 + \sigma_1^2)\boldsymbol{I} + Nh_I^2\boldsymbol{D}/M)$

For single-entry transformed Z score, $\tilde{Z}_J \sim N(0, Nh_I^2D_{jj}/M + 1 + 2(h_I^2 + \sigma_1^2))$

Finally, we derive that $Var(\tilde{Z}_J) = 2\left(Nh_I^2D_{jj}/M + 1 + 2(h_I^2 + \sigma_1^2)\right)^2$

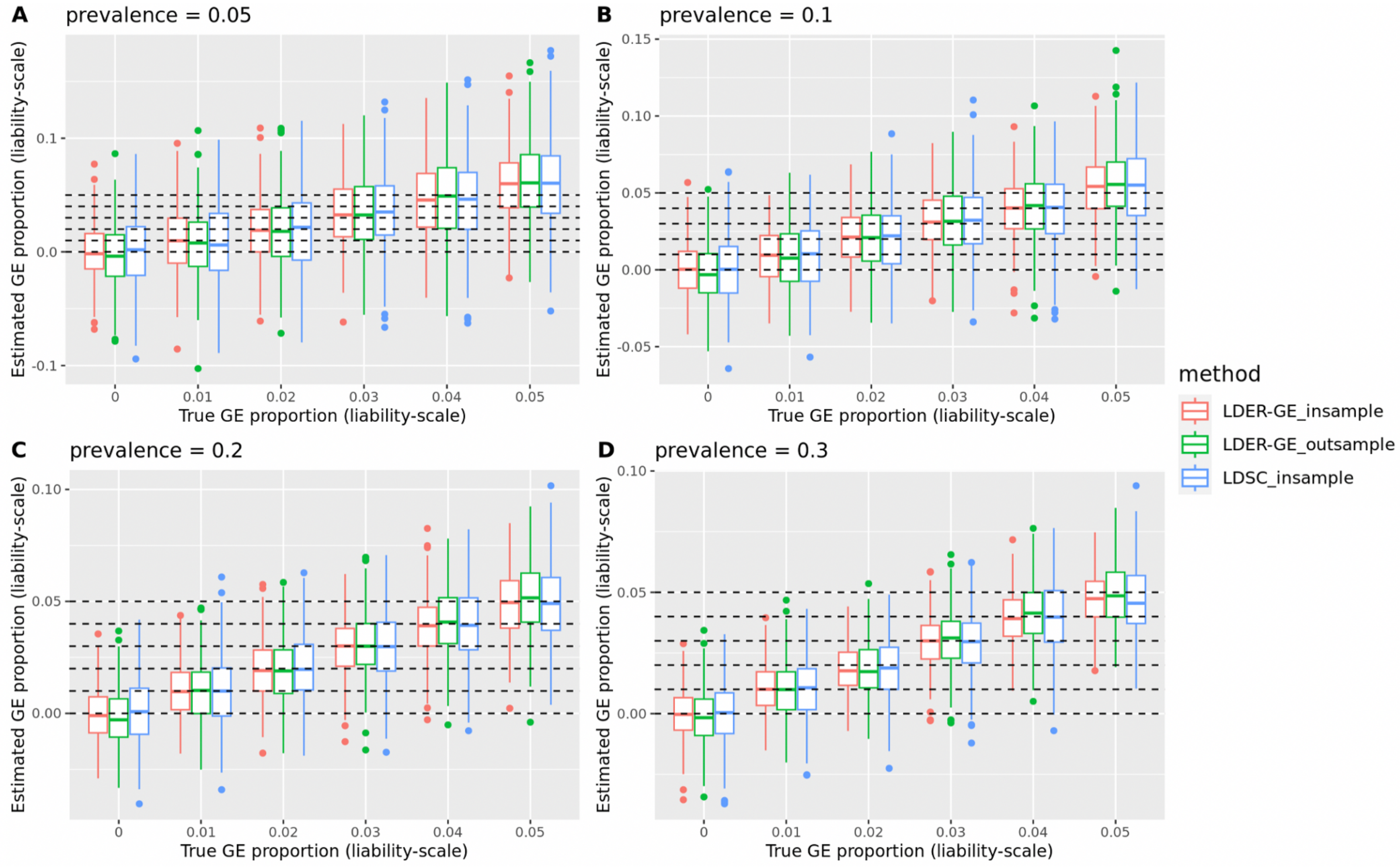

Supplementary Figure 1: Histogram comparison of LDSC-based method and LDER-GE with in-sample and out-sample reference panel on simulations from real genotype panel, binary phenotype.

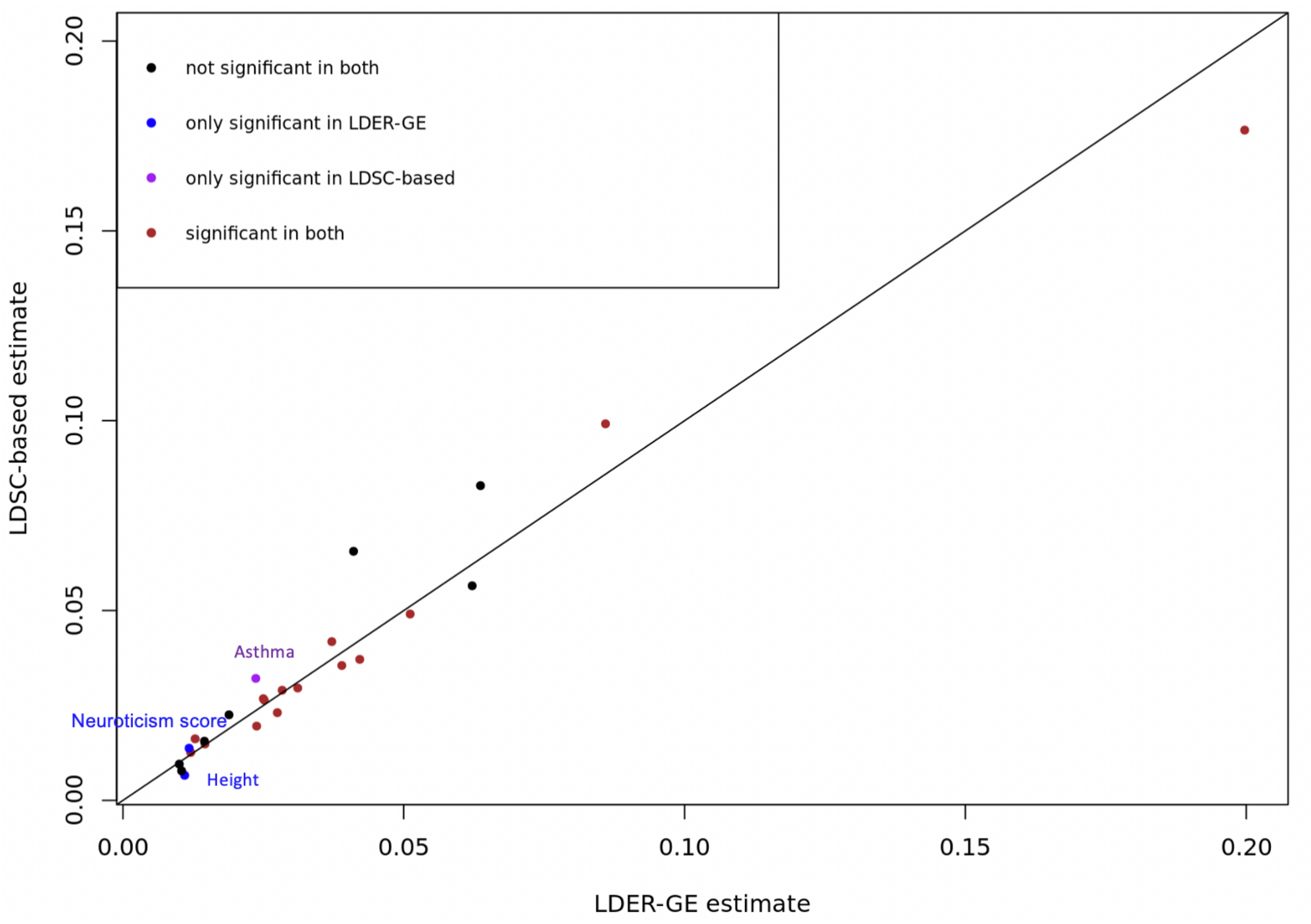

Supplementary Figure 2: Multi-GE interaction variance estimates from LDSC-based method and LDER-GE of 25 phenotypes in UKBB dataset. For binary phenotypes, GE interaction variance is reported on the liability-scale.
